# Homogenization of taxonomic, phylogenetic, and functional characteristics of bumble bee communities at regional scales in anthropogenic landscapes

**DOI:** 10.1101/2023.10.13.562048

**Authors:** Kayla I Perry, Claudio Gratton, Taylor Tai, James P Strange

## Abstract

Biotic homogenization has been documented following extensive anthropogenic landscape change such as urbanization and agriculture, but diverse native communities also have been reported in these ecosystems. Understanding the influence of landscape-level characteristics on processes of community assembly can inform how human-dominated landscapes shape the structure and composition of local communities, including important pollinators such as bumble bees (*Bombus* spp.). The objective of this study was to investigate multi-scale patterns of taxonomic, phylogenetic, and functional beta-diversity of bumble bees in greenspaces along an urban-agricultural gradient to understand landscape-scale constraints on processes of community assembly. Bumble bees were collected in greenspaces along an urban-agricultural gradient in Madison, WI, USA. Patterns of biotic homogenization were investigated using measures of beta-diversity and null models relative to a regional bumble bee species pool in a 100 km area surrounding the city. Nine of the expected 13 species from the regional pool were collected in greenspaces in urban and agricultural landscapes. At the regional scale, we found evidence of taxonomic, phylogenetic, and functional homogenization among bumble bee communities in urban and agricultural landscapes, with species that were smaller in size, had shorter wings, were less hairy, but had larger eyes and longer setae on the corbicula (pollen-carrying hind legs) being more common than expected based on null models. When we evaluated filtering from the anthropogenic species pools (i.e., urban and agricultural) to local greenspaces, we found nuanced differences among land cover types, wherein agricultural landscapes supported higher beta-diversity of bumble bee communities than expected while urban landscapes continued to show signals of homogenization. Overall, anthropogenic landscapes acted as a strong filter for bumble bees, broadly selecting for a subset of functionally similar and phylogenetically related species that resulted in homogenization of communities within the region. Our findings support a landscape-level approach to biodiversity conservation that promotes diversifying landscapes to support diverse pollinator populations.

**OPEN RESEARCH STATEMENT:** Data and novel code associated with this submission are provided in an external repository to be evaluated during the peer review process and are available at https://github.com/kiperry/WI_Bumble_Bees. If this paper is accepted for publication, data and code will be permanently archived within a linked Zenodo repository.

## INTRODUCTION

Human-mediated landscape change is a major driver of natural habitat loss and fragmentation (McKinney 2002, Grimm et al. 2008), with consequences for biodiversity and ecosystem functioning (Fischer and Lindenmayer 2007, Cardinale et al. 2012, Haddad et al. 2015). Anthropogenic conversion of natural habitat to urban or agricultural landscapes has occurred on a global scale (Foley et al. 2005, Seto et al. 2011, Fragkias et al. 2013), altering patterns of species diversity and ecosystem services (Dirzo and Raven 2003, Hooper et al. 2012, Dirzo et al. 2014).

Anthropogenic modification of natural habitat can lead to losses of local species diversity (i.e., alpha-diversity) as well as an increase in the compositional similarity (i.e., lower beta-diversity) of communities between locations (McKinney and Lockwood 1999, Gossner et al. 2016). Biotic homogenization, or the increased similarity of communities due to shared traits of species that allow them to persist in anthropogenically altered environments, is predicted in landscapes altered by human activities such as urbanization (McKinney 2006, Knop 2016) and agricultural intensification (Ekroos et al. 2010, Karp et al. 2012, Gámez-Virués et al. 2015) due to environmental filtering processes. For example, homogenization of bird communities has been documented with increasing urbanization intensity wherein cities had fewer habitat specialists and ground nesting species (Clergeau et al. 2006, Luck and Smallbone 2011). Because the composition of species within and among communities relates to their roles in ecosystem services and function (Mace et al. 2012), regional biotic homogenization may reduce ecosystem multifunctionality (i.e., carbon sequestration, pollination, food production, nutrient cycling, pest suppression, water purification) at the landscape-scale (van der Plas et al. 2016) and should be a priority for biodiversity conservation management.

To support diverse communities within human-dominated landscapes, there is a need to understand the drivers of processes of community assembly. Community assembly is determined by a series of hierarchical abiotic and biotic filters operating at various spatial scales that cumulatively influence which species from the regional species pool can colonize and establish to form local communities (Fig. 1) (HilleRisLambers et al. 2012, Aronson et al. 2016, Perry et al. 2020). At the regional scale, broad geographic and climatic factors as well as past and current anthropogenic land use practices that determine the composition, configuration, and connectivity of natural habitat can influence the regional species pool and thus, the species that can be potentially found within local communities.

**Figure 1.**
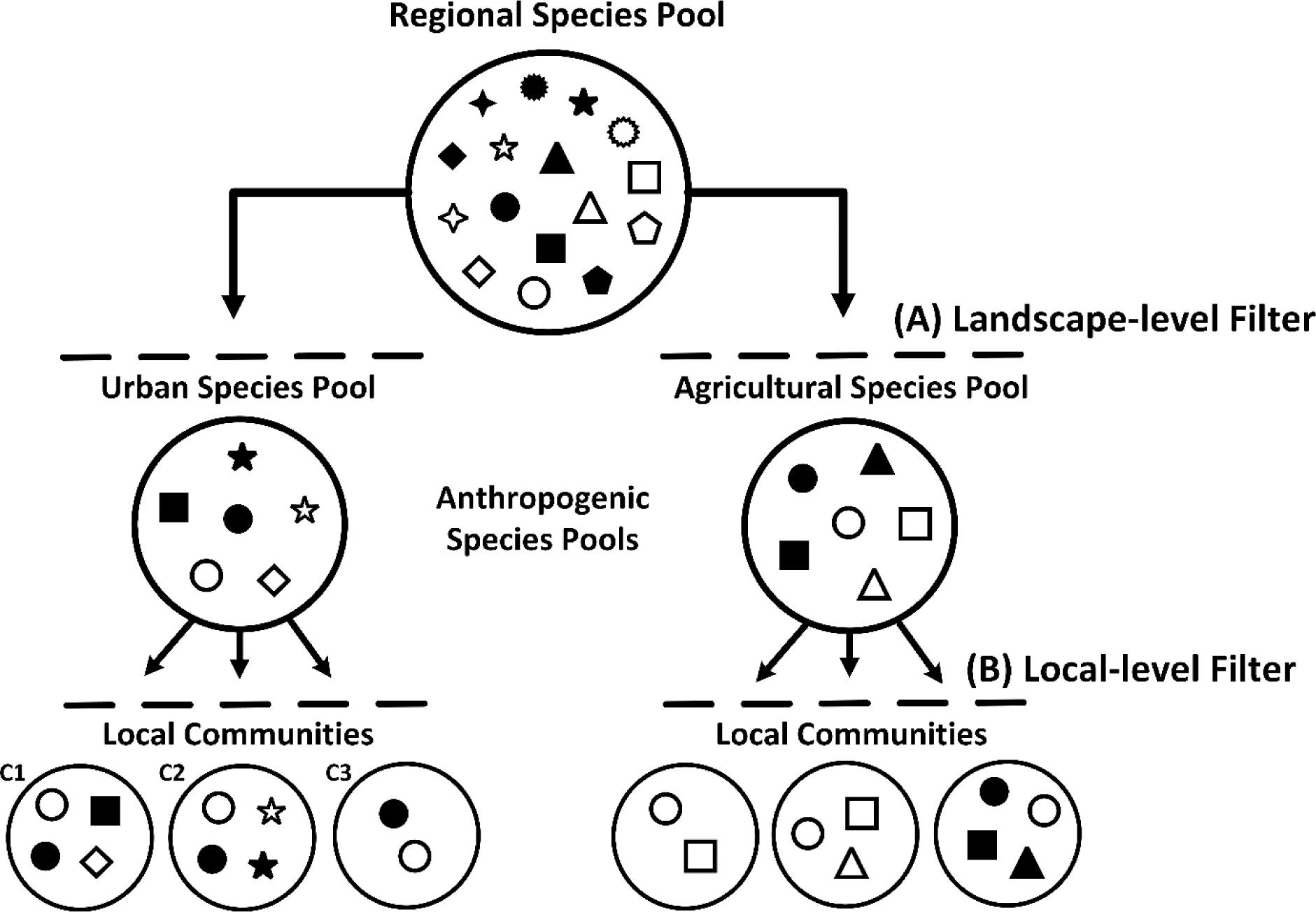
Conceptual model of community assembly in human-dominated landscapes. The regional species pool (gamma-diversity) represents all species within a given area that have the opportunity to occur within local communities (alpha-diversity). In human-dominated landscapes, anthropogenic filters are predicted to create additional secondary species pools (i.e., urban or agricultural) which have the capacity to persist due to features of the habitats such as disturbance regimes (e.g., pesticide use, tillage), biotic (e.g, resource availability), or abiotic (e.g., temperature, moisture) factors. Assembly of local communities is influenced by a combination of (A) landscape-level processes such as species’ dispersal capacity, immigration-emigration dynamics, and chance events that influence colonization success and (B) local filters associated with among site variation in species interactions, resource availability, and the environment influence establishment success. Beta-diversity is a measure of compositional similarity among local communities, which can be decomposed into processes of turnover (reflects replacement) and nestedness (reflects loss or gain). High turnover is represented by C1 and C2, as two species are shared between these local communities while two species are replaced from the urban species pool. High nestedness is represented by C1 and C3, as two species are lost making the species present in C3 a subset of those in C1. Symbols of different shapes and colors represent different species.

Landscapes that have been highly modified by humans are predicted to have species pools that are created by anthropogenic filters acting on the regional pool (Fig. 1A: Landscape-level Filter) (Aronson et al. 2016). For example, urban landscapes are dominated by built infrastructure and impervious surfaces that may act as barriers for some species (Banaszak-Cibicka and Żmihorski 2012, Jha and Kremen 2013, Perry et al. 2020), creating a secondary urban species pool (Fig. 1A, left). In this case, the observed urban pool consists of species that have been filtered from the regional pool because they are able to successfully colonize and establish within highly urbanized environments. These landscape-level anthropogenic filters are predicted to form a secondary species pool by selecting for a subset of generalist, functionally and phylogenetically similar species from the regional pool that are well-adapted to such human-dominated environments. For example, homogenization (i.e., lower beta-diversity) of butterfly and diurnal moth communities was observed in landscapes with high agricultural intensity, and these patterns were driven by an increase in habitat generalist and good disperser species (Ekroos et al. 2010). Subsequently, local communities are formed by species represented within the secondary species pool (Fig. 1B: Local-level Filter), which have the capacity to persist due to features of the habitats such as disturbance regimes (e.g., pesticide use, tillage), biotic (e.g., resource availability) or abiotic (e.g., temperature, moisture) factors.

Effects of anthropogenic landscapes on community assembly can be assessed using measures of beta-diversity. Because beta-diversity links spatial patterns of diversity at local scales (alpha-diversity) with the regional species pool (gamma-diversity), this diversity metric can be used to investigate patterns of biotic homogenization (Anderson et al. 2011, Cornell and Harrison 2014). Shifts in beta-diversity among communities can reflect two different processes, turnover and nestedness (Baselga 2010). Spatial turnover among communities indicates the replacement of some species by others (Fig. 1, C1 & C2), while spatial nestedness indicates the non-random loss or gain of species such that some communities are a subset of others (Fig. 1, C1 & C3). Patterns of beta-diversity that show low species turnover and high species nestedness suggest an increase in the compositional similarity among communities and are indicative of homogenization (Olden et al. 2004, Olden and Rooney 2006). Based on this biotic homogenization hypothesis, we predict that for mobile insects occurring across a diverse urban-agricultural mosaic anthropogenic land use practices and legacies will result in lower alpha- and beta-diversity of communities.

Pollinators include highly mobile insects that provide essential ecosystem services which support wild plants and cultivated crops (Klein et al. 2007, Ollerton et al. 2011), including in urban agricultural systems (Matteson and Langellotto 2009, Potter and LeBuhn 2015). Declines in wild and domesticated pollinator populations have been documented globally with consequences for the resilience of pollination services (Potts et al. 2010). Pollinators experience many environmental stressors at landscape and local scales (Gill et al. 2016). Of those stressors, anthropogenic land use change and legacies that result in the loss and fragmentation of natural habitat are hypothesized as important drivers of pollinator declines (Potts et al. 2010, Vanbergen et al. 2013). However, while urban and agricultural landscapes result in the loss and fragmentation of natural habitat, these ecosystems can be highly heterogeneous in resource and nesting availability with context dependent impacts for pollinator populations. For example, impacts of urbanization on pollinators have been inconsistent, and studies have documented positive, neutral, and negative relationships (Hernandez et al. 2009, Ahrné et al. 2009, Baldock et al. 2019, Wenzel et al. 2020, Fournier et al. 2020, Theodorou et al. 2020) possibly due to high heterogeneity in the characteristics of urban landscapes. A more mechanistic approach is required to explain observed biodiversity patterns and to understand the relative importance of anthropogenic landscape-scale processes as drivers of pollinator community assembly.

Functional diversity, captured as the value and range of species’ traits, provides a mechanistic means for understanding patterns of biodiversity and processes of community assembly (McGill et al. 2006, Mason and de Bello 2013). Functional traits include any morphological, physiological, phenological, or behavioral character that is measurable and impacts the fitness of an individual either directly or indirectly (Violle et al. 2007). Functional trait-based approaches link the taxonomic composition of communities with the ecological and life history strategies of species to understand the processes that govern community assembly under various environmental conditions (De Bello et al. 2021). Therefore, functional diversity is considered a better predictor of ecosystem function than more traditional taxonomic diversity metrics (Gagic et al. 2015). From a community assembly perspective, a functional diversity approach can be used to understand the traits that are selected for or against in anthropogenic landscapes, providing insight into why certain species are more successful in these environments than others. Phylogenetic diversity, a representation of the evolutionary distances among species, provides an additional dimension of diversity based on the relatedness among species. Because many related species share similar traits, functional and phylogenetic diversity metrics are often used together to understand the influence of community changes on ecosystem function (Srivastava et al. 2012). Incorporating a functional trait-based approach with more traditional taxonomic and phylogenetic diversity metrics can provide a comprehensive assessment of pollinator communities in anthropogenic landscapes.

Bumble bees (*Bombus* spp.) are an important group of insect pollinators that have experienced population declines and range reductions worldwide (Grixti et al. 2009, Williams et al. 2009, Cameron et al. 2011, Colla et al. 2012). Although the causes of decline are likely multifaceted (Goulson et al. 2015), human-mediated landscape change such as urbanization and agricultural intensification are predicted to be key drivers (Grixti et al. 2009, Cameron et al. 2011, Hemberger et al. 2021). Studies have found that bumble bee species are differentially impacted by stressors such as urbanization, with declines reported for some species while others remain abundant and widespread (Ahrné et al. 2009, Banaszak-Cibicka and Żmihorski 2012). For example, declines in captures of 11 bumble bee species native to eastern North America were documented via museum records over the past century, while captures of eight species have remained stable or increased over this same time period (Colla et al. 2012). Populations of *Bombus impatiens* Cresson have remained widespread in the United States, while populations of *Bombus affinis* Cresson have declined in geographic distribution from its historic range (Cameron et al. 2011), suggesting differences in how species respond to anthropogenically altered habitats (Hemberger et al. 2021). Differential responses of bumble bee species to anthropogenic stressors may be explained by their life history traits such as those related to dispersal capacity, nesting preferences, diet, and phenology. For example, shifts in body size of bumble bees have been associated with urbanization (Austin et al. 2022), with the direction of the response (positive or negative) being species-specific (Eggenberger et al. 2019, Theodorou et al. 2021). Body size is a key trait linked to functions such as dispersal capacity, metabolism, and reproduction (Moretti et al. 2017). If some life history traits are selected for or against in response to anthropogenic landscape-level filters, declines in some bumble bee species but not others also may increase the functional similarity among communities leading to functional homogenization (Smart et al. 2006). Whether different anthropogenic landscapes (i.e., urban vs. agricultural) select for a similar or divergent subset of life history traits is not well understood but could impact the maintenance of diversity and ecosystem services at large spatial scales.

To understand whether anthropogenic landscapes dominated by urban or agricultural land cover result in biotic homogenization of pollinator communities, this study investigated multi-scale processes of bumble bee community assembly along an urban-agricultural gradient (Fig. 1A, B). To do this, we evaluated patterns of taxonomic, phylogenetic, and functional beta-diversity of bumble bees among greenspaces via null model analyses of regional, anthropogenic, and local species pools. If processes of community assembly differ based on the dominant land cover type (i.e., urban or agricultural), we hypothesize that a different subset of bumble bee species with different traits will be filtered from the regional species pool to create the urban and agricultural species pools (Fig. 1A). We specifically hypothesize that: 1) taxonomic, phylogenetic, and functional beta-diversity of bumble bees will be more similar among greenspaces in urban and/or agricultural landscapes than expected by chance, via null model analysis based on the regional species pool; and 2) greenspaces in urban and/or agricultural landscapes will have only a subset of phylogenetically related bumble bee species with similar traits from the regional species pool, resulting in patterns of taxonomic, phylogenetic, and functional beta-diversity driven by high nestedness and low turnover. Within anthropogenic species pools (i.e., urban and agricultural), we hypothesize that among site variation in abiotic and biotic factors will select for different subsets of bumble bee species with different traits to create local communities (Fig. 1B).

## METHODS

### Study area

This study was conducted along an urban-agricultural gradient in Dane County, Wisconsin, USA, where the city of Madison is located. Sites were selected through a multi-step grid-based procedure wherein Dane County was divided into a 1 x 1 km array to limit any landscape-level confounding factors such as geographic location or land cover characteristics. The predominant land cover class (i.e., urban or agriculture) was determined for each 1 x 1 km grid cell, and 25 1 km grid cells were randomly selected from each land cover class using a spatially stratified approach. Each of those 1 km grid cells was further divided into 100 x 100 m blocks, and five 50 m radius circles were randomly placed within each block.

The first three 50 m radius circles that were accessible were selected as survey sites. In 2019, 47 survey sites across 16 blocks were established. An additional 12 survey sites across four blocks were established in 2020, totaling 20 sampling blocks. Additional information on site selection is provided by Nunes et al. (in revision).

### Bumble bee collection and identification

At each survey site, a 100 m transect was established based on the availability of floral resources for bumble bees and location accessibility, hereafter referred to as “greenspaces”. Bumble bee observations were conducted along transects where a surveyor walked a zig-zag pattern changing direction every 10 m, with each short segment taking ∼2 min, and with pauses of 1 min at each direction change (total ∼30 min per 100 m transect). Bumble bees observed along the transects were collected and identified to species, stopping the timer during handling of bees. Any individuals of the federally endangered *B. affinis* were recorded and immediately released, per handling requirements of the U.S. Fish and Wildlife Service Permit.

Transects were surveyed for bumble bees three times in 2019 (6 June – 7 July; 8 July – 31 July; 2 August – 20 August) and 2020 (8-19 June; 22 June – 28 July; 8-21 August). University imposed personnel restrictions in 2020 in response to the SARS-CoV-2 pandemic resulted in reduced sampling frequency of some transects previously established in 2019. Additional details about bumble bee sampling protocols are provided by Nunes et al. (X).

### Regional species pool

The regional species pool was characterized within a 100 km radius surrounding Madison, Wisconsin using bumble bee species records observed from 2017-2021 from the citizen science project Bumble Bee Watch (https://www.bumblebeewatch.org/) and the literature (Wolf and Ascher 2009).

Twenty-seven counties in Wisconsin and Illinois fall within the 100 km area surrounding the city. Bumble bee species were included in the regional pool if verified records were reported from any of the 27 counties. This list of species was cross-referenced with the list provided in Wolf and Ascher (2009), where the counties overlapped with the southwestern, southeastern, and central sands landscape regions. To reduce potential bias associated with sample collection times and potential for observations of kleptoparasitic species, only species with verified records in Bumble Bee Watch were included in the regional pool, which generated a conservative list of 13 bumble bee species (Table 1).

**Table 1.** Bumble bee species identified within the regional pool and abundances of those species collected in greenspaces embedded in urban and agricultural landscapes surrounding Madison, Wisconsin, USA in 2019 and 2020.

| Species | Urban | Agriculture | Total |
| --- | --- | --- | --- |
| *Bombus impatiens* Cresson | 644 | 192 | 836 |
| *Bombus vagans* Smith | 153 | 94 | 247 |
| *Bombus bimaculatus* Cresson | 114 | 44 | 158 |
| *Bombus griseocollis* (DeGeer) | 76 | 29 | 105 |
| *Bombus rufocinctus* Cresson | 62 | 37 | 99 |
| *Bombus fervidus* (Fabricius) | 22 | 10 | 32 |
| *Bombus auricomus* (Robertson) | 3 | 21 | 24 |
| *Bombus affinis* Cresson | 11 | 2 | 13 |
| *Bombus borealis* Kirby | 3 | 4 | 7 |
| *Bombus citrinus* (Smith) | 0 | 0 | 0 |
| *Bombus pensylvanicus* (DeGeer) | 0 | 0 | 0 |
| *Bombus perplexus* Cresson | 0 | 0 | 0 |
| *Bombus ternarius* Say | 0 | 0 | 0 |
| **Total** | **1088** | **433** | **1521** |

### Trait and phylogenetic data

Seventeen morphological and ecological traits were selected for this study because they are considered important for species’ responses to the environment across various spatial scales (Moretti et al. 2017). Five traits were determined for each species from the literature (Williams et al. 2014; DiscoverLife.org). These were tongue length (short, medium, or long), nest location (primarily aboveground, belowground, or kleptoparasitic), and body length range for queen, worker, and male castes. Body length range was calculated by taking the difference between the low and high values reported for the size range of each species. An additional twelve traits were measured from at least five workers of each species (range 5-12 individuals of each species). Traits were measured under a dissecting microscope using an eyepiece micrometer (reticle) to the nearest 0.1 mm and included: inter-tegular distance, wing marginal cell length, wing length and width, head width, eye length and width, corbicula length and width, scape length, length of setae on the posterior edge of the corbicula, and length of hair on the thorax. To control for variation in bee body size, relative measurements of wing length and width, head width, eye length and width, scape length, and corbicula length and width were calculated as their ratio to inter-tegular distance for each individual. Trait measurements were averaged across individuals to calculate species-specific means.

Phylogenetic relationships among bumble bee species are based on the phylogeny developed by Cameron et al. (2007). The phylogeny was trimmed using the drop.tip function in the R package ‘ape’ (Paradis and Schliep 2019) to produce a phylogenetic tree that included only the 13 species determined to be in the regional species pool.

### Beta-diversity metrics

All beta-diversity metrics are based on presence/absence data of bumble bee species occurrences among sites. The occurrences of bumble bee species were pooled across sites within a block, and then pooled across years. Bumble bee communities were assessed using multiple beta-diversity metrics: taxonomic (traditional approach based on the presence/absence of species among sites), phylogenetic (similar to the taxonomic approach but weighted by species relatedness), and functional (approach based on species traits among sites). Patterns of observed bumble bee taxonomic, phylogenetic, and functional beta-diversity were evaluated by calculating pairwise Sorensen dissimilarity (β_sor_) matrices among sites, and then decomposing total beta-diversity into a turnover component (β_sim_; reflects replacement) and a nestedness component (β_sne_; reflects loss or gain) (i.e., β_sor_ = β_sim_ + β_sne_) (Baselga 2010). These matrices were generated using the R package ‘betapart’ (Baselga and Orme 2012). Phylogenetic beta-diversity is based on Faith’s PD index (Faith 1992) which uses the sum of the branch lengths of the tree as a representation of evolutionary distances to construct the pairwise distance matrix.

For functional beta-diversity, a trait distance matrix was created using the R package ‘gawdis’ (De Bello et al. 2021), which allows for continuous and categorical traits. Traits were weighted with the optimized argument that uses 300 iterations to identify equal contributions of each trait to the distance matrix. Additionally, traits on the head (i.e., head width and eye length) were grouped together to limit their combined contribution, as these traits were more correlated to each other than other morphological traits on the body. Next, a principal coordinates analysis used the trait distance matrix to generate condensed trait axes via the R package ‘ade4’ (Dray and Dufour 2007). Because one site had only three observed species in the community, we were only able to use two trait axes for the functional beta-diversity metrics. To further explore patterns of bumble bee functional diversity among sites, observed community-weighted means (CWMs) for all continuous and categorical traits were calculated using the R package ‘FD’ (Laliberté et al. 2014). This metric represents the average value of a specific trait for a local community (i.e., at a site) weighted by the relative abundance of the observed species.

### Landscape data

Landscape data were obtained from the 2016 National Land Cover Database (https://www.mrlc.gov/data/nlcd-2016-land-cover-conus), which is a comprehensive raster data layer generated for the continental United States at a 30 m resolution. For each site, circular buffers with 1.5 km radii were created using ArcGIS v10.8.1 (ESRI 2020). This spatial scale was selected because it corresponds to foraging distances of bumble bees (Osborne et al. 2008). Fifteen land cover classes were represented across these buffers: Open Water, Developed/Open Space, Developed/Low Intensity, Developed/Medium Intensity, Developed/High Intensity, Barren Land, Deciduous Forest, Evergreen Forest, Mixed Forest, Shrub/Scrub, Herbaceous, Hay/Pasture, Cultivated Crops, Woody Wetlands, and Emergent Herbaceous Wetlands.

Using these data, we calculated metrics that quantify landscape composition, configuration, and connectivity using the software program FRAGSTATS (McGarigal and Marks 1994). All metrics were calculated based on the 8-neighbor rule for delineating patches where the eight adjacent cells were considered members of the same patch. Landscape composition was quantified by measuring percentage of urban land cover (sum of Developed/Low Intensity, Developed/Medium Intensity, Developed/High Intensity), percentage of agricultural land cover (sum of Hay/Pasture, Cultivated Crops), and percentage of natural habitat (sum of Deciduous Forest, Evergreen Forest, Mixed Forest, Shrub/Scrub, Herbaceous, Woody Wetlands, and Emergent Herbaceous Wetlands). Based on these percentages, all sites were classified as dominated (i.e., >50% of land cover area) by either urban (12 sites) or agricultural (8 sites) land cover. Because the primary natural land cover type surrounding our sites was forest, landscape configuration and connectivity were measured using the metrics Largest Patch Index (percentage of the landscape comprised by the largest patch of a given land cover type), Edge Density (length of edge, m/ha), and Euclidean Nearest Neighbor Distance (average distance to the nearest neighboring patch of the same land cover type, m) for class forest (Deciduous Forest, Evergreen Forest, Mixed Forest). Simpson’s Patch Diversity Index (measures diversity of patch types) was calculated based on 9 land cover classes: Open Water, Urban Greenspace (Developed/Open Space), Urban Developed (Developed/Low Intensity, Developed/Medium Intensity, Developed/High Intensity), Barren Land, Forest (Deciduous Forest, Evergreen Forest, Mixed Forest), Shrubland (Shrub/Scrub), Grassland (Herbaceous), Agriculture (Hay/Pasture, Cultivated Crops), and Wetland (Woody Wetlands, Emergent Herbaceous Wetlands).

### Statistical analysis

Pearson correlation analyses were used to assess the relationships among bumble bee traits. The traits wing marginal cell length, wing width, eye width, corbicula width, and scape length had correlation coefficients of 0.60 or higher with other traits and were removed from further analyses (see Appendix S1: Figure S1). Traits were checked for normality and log transformed if necessary. All analyses were conducted in R 4.0.3 (R Core Team 2020). All data and code for the analyses are available at: https://github.com/kiperry/WI_Bumble_Bees.

Null models were developed to investigate processes of bumble bee community assembly at two scales: 1) from the regional species pool to the anthropogenic (i.e., urban and agricultural) species pools; and 2) from the anthropogenic species pools to local greenspaces. For each scale, null models randomized the species presence/absence matrix, trait matrix, and phylogenetic tree for each of 999 iterations.

Standardized effect sizes (SES) were calculated for each metric to determine if observed values were higher or lower than expected by chance based on the randomized communities. Wilcoxon rank-sum tests were used to assess whether SES values differed from null model expectations.

To assess whether processes of community assembly differed based on the dominant land cover type (i.e., urban or agricultural), we developed a null model to evaluate filtering from the regional species pool to the anthropogenic species pools (Fig. 1A). In this null model, the species presence/absence matrix, trait matrix, and phylogenetic tree included data for all 13 species determined to be in the regional species pool. The presence/absence species matrix was randomized using the randomizeMatrix function in the R package ‘picante’ (Kembel et al. 2010) with the richness null model implemented that maintains the same sample species richness. Because this dataset included species determined to be in the regional pool but which were not collected at any of the sites (i.e., an occurrence of 0 across all sites; *Bombus citrinus* (Smith), *Bombus pensylvanicus* (DeGeer), *Bombus perplexus* Cresson, and *Bombus ternarius* Say), this null model did not maintain species occurrence frequency with each randomization. The trait matrix was randomized by shuffling rows of traits among species, which maintains patterns of covariation among traits. Species relationships on the phylogenetic tree were randomized using the function tipShuffle in the R package ‘picante’ (Kembel et al. 2010). Metrics included in the regional species pool to anthropogenic species pools null model were the CWMs for each trait and measures of taxonomic, phylogenetic, and functional beta-diversity (total beta-diversity and the turnover and nestedness components).

To assess whether processes of community assembly differed based on variation among sites embedded in either urban-dominated or agricultural-dominated landscapes, we developed null models to evaluate filtering from the anthropogenic species pools to local greenspaces (Fig. 1B). In these null models, the species presence/absence matrix, trait matrix, and phylogenetic tree included data for only those species collected during the study to represent the secondary anthropogenic species pools. The presence/absence species matrix was randomized using the randomizeMatrix function with the independent swap null model implemented which maintains sample species richness and species occurrence frequency (i.e., common species remain common and rare species remain rare). Similar to the null model above, the trait matrix was randomized by shuffling rows of traits among species and species were randomized on the phylogenetic tree using the tipShuffle function. Metrics included in the anthropogenic species pools to local greenspaces null models were the CWMs for each trait and measures of taxonomic, phylogenetic, and functional beta-diversity (total beta-diversity and the turnover and nestedness components).

Partial least squares canonical analysis (PLSCA) and relevance network analysis were used to further investigate the importance of variation in landscape characteristics, particularly the composition, configuration, and connectivity of natural habitat, for the assembly of bumble bee communities among local greenspaces. Therefore, these analyses assessed relationships among SES values for bumble bee CWM metrics calculated based on species observed in the anthropogenic species pools and landscape variables. This technique analyzes the linear relationships among variables in two matrices and maximizes the covariance explained between them by deriving a latent variable from each matrix (Abdi and Williams 2013). Partial least squares methods have several advantages which include the ability to incorporate multiple response variables, use many predictors that may be collinear, and have small sample sizes (Carrascal et al. 2009). In the PLSCA, the following landscape variables were included: 1) landscape patch diversity; 2) percentage agriculture; 3) percentage urban; 4) percentage natural habitat; 5) forest largest patch index (LPI Forest); 6) forest edge density (ED Forest); and 7) forest Euclidean nearest neighbor distance (ENN Forest). The PLSCA was initially conducted with the complete set of CWM metrics calculated for bumble bees. Variables were scaled to have a mean of zero and variance of one. Following these initial analyses, bumble bee CWMs and landscape variables with PLSCA loadings <0.30 were removed (see Appendix S1: Table S1), and an additional PLSCA and relevance network analyses were conducted with reduced datasets. Relevance network analysis calculates pairwise similarity values which approximate a Pearson correlation. Similarity values are calculated by summing the correlations between individual variables and each of the latent variables from the PLCA. A 0.5 similarity value threshold was used to assess the strength of the variable associations, within similarity values higher than 0.5 or lower than -0.5 considered significant. PLSCA and relevance network analysis were performed using the package ‘mixOmics’ (Rohart et al. 2017) in R.

## RESULTS

A total of 1,521 bumble bees representing nine species were collected during 2019-2020, which comprised 69% of species identified within the regional species pool (Table 1). Of the nine species observed, *Bombus impatiens* Cresson and *Bombus vagans* Smith had the highest occurrences as these two species were collected in all survey blocks, followed by *Bombus bimaculatus* Cresson (observed in 95% of survey blocks), *Bombus griseocollis* (DeGeer) (90%), and *Bombus rufocinctus* Cresson (70%).

Thirteen individuals of *Bombus affinis* Cresson were observed during this study. All nine species were observed in greenspaces in urban and agricultural landscapes (Table 1).

### Regional species pool to anthropogenic species pools

The observed total taxonomic, phylogenetic, and functional beta-diversity metrics were lower than null model expectations based on the regional species pool, and these patterns were the same for urban and agricultural landscapes, suggesting a strong landscape-level filter (Fig. 2; Table 2A). Bumble bee communities sampled in urban and agricultural landscapes contained a similar subset of species that were more phylogenetically related than expected. These species were characterized by a similar subset of traits represented in the regional species pool such that bumble bee communities also were more functionally similar than expected. Patterns of total taxonomic, phylogenetic, and functional beta-diversity were driven by processes related to turnover and nestedness. Taxonomic, phylogenetic, and functional turnover were lower than null model expectations in urban and agricultural landscapes (Fig. 2; Table 2A), indicating lower levels of species and trait turnover among bumble bee communities.

**Figure 2.**
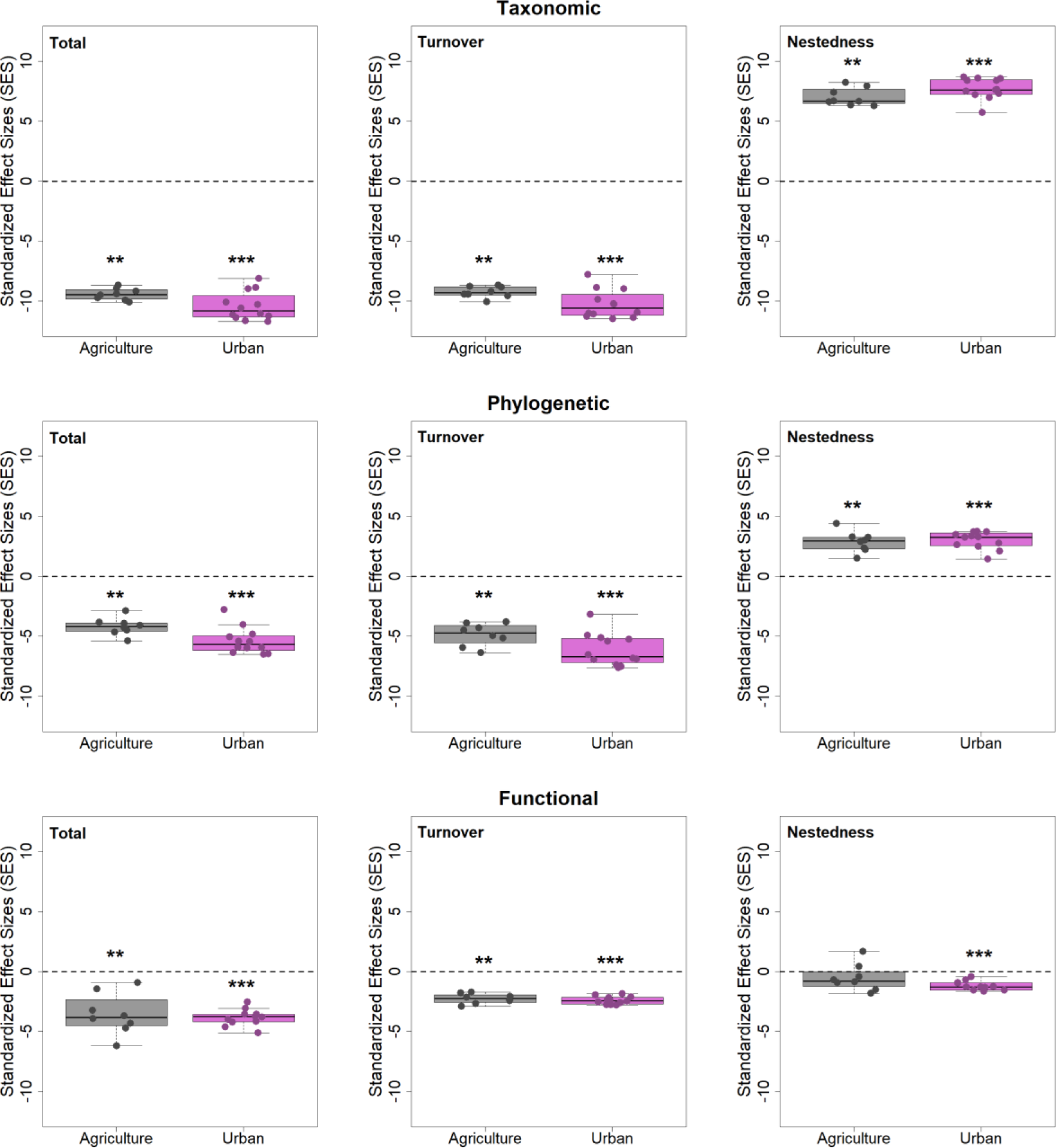
Standardized effect sizes (SES) of observed bumble bee taxonomic (top), phylogenetic (middle), and functional (bottom) beta-diversity in greenspaces embedded in agricultural (left) and urban (right) landscapes surrounding Madison, Wisconsin, USA. Observed values were compared to null model expectations based on the regional pool of 13 species (Fig. 1A, Landscape-level Filter). Positive SES values indicate observed beta-diversity metrics were higher than null model expectations, while negative SES values indicate observed beta-diversity metrics were lower than null model expectations. Pairwise Sorensen dissimilarity (β_sor_) matrices were calculated for taxonomic, phylogenetic, and functional beta- diversity. Total beta-diversity for each metric was decomposed into a turnover component (β_sim_; reflects replacement) and a nestedness component (β_sne_; reflects loss or gain) (i.e., β_sor_ = β_sim_ + β_sne_). Wilcoxon rank sum tests were used to assess whether SES values differed from null model expectations. * P<0.05, ** P<0.01, ***P<0.001, otherwise results were not significant at P>0.05.

**Table 2.** Standardized effect sizes (SES) of observed bumble bee taxonomic, phylogenetic, and functional beta-diversity as well as community- weighted means for each trait in greenspaces embedded in urban and agricultural landscapes surrounding Madison, Wisconsin, USA. Observed values were compared to null model expectations based on the regional pool of 13 species (Fig. 1A, Landscape-level Filter). Positive SES values indicate observed metrics were higher than null model expectations, while negative SES values indicate observed metrics were lower than null model expectations. Wilcoxon rank sum tests were used to assess whether SES values differed from null model expectations. * P<0.05, ** P<0.01, ***P<0.001, otherwise results were not significant at P>0.05.

| Metric | Category/Index | Agriculture |  | Urban |  |
| --- | --- | --- | --- | --- | --- |
|  |  | SES<br>(pseudo)median | 95% CI | SES<br>(pseudo)median | 95% CI |
| A) |  |  |  |  |  |
| Taxonomic beta-diversity | Total | -9.444** | -9.922, -8.948 | -10.585*** | -11.266, -9.632 |
|  | Turnover | -9.209** | -9.748, -8.777 | -10.334*** | -11.157, -9.452 |
|  | Nestedness | 7.021** | 6.446, 7.827 | 7.818*** | 7.159, 8.383 |
| Phylogenetic beta-diversity | Total | -4.216** | -4.843, -3.483 | -5.567*** | -6.165, -4.632 |
|  | Turnover | -4.852** | -5.763, -4.051 | -6.218*** | -7.148, -5.181 |
|  | Nestedness | 2.855** | 2.200, 3.697 | 3.056*** | 2.507, 3.497 |
| Functional beta-diversity | Total | -3.656** | -4.881, -2.075 | -3.747*** | -4.149, -3.305 |
|  | Turnover | -2.253** | -2.673, -1.931 | -2.422*** | -2.632, -2.191 |
|  | Nestedness | -0.543 | -1.366, 0.595 | -1.182*** | -1.392, -0.850 |
| B) |  |  |  |  |  |
| Body length | Inter-tegular distance | -0.535* | -1.170, -0.016 | -0.764*** | -1.062, -0.507 |
| Body length variance | Queen | 0.099 | -0.558, 0.607 | -0.021 | -0.501, 0.316 |
|  | Male | -0.016 | -0.711, 0.690 | -0.094 | -0.201, 0.158 |
|  | Worker | 0.431 | -0.256, 1.083 | 0.717*** | 0.424, 1.181 |
| Tongue length | Short | -0.294 | -1.087, 0.608 | 0.356* | 0.146, 0.583 |
|  | Medium | 0.168 | -0.392, 0.758 | 0.214 | -0.006, 0.502 |
|  | Long | 0.071 | -0.588, 0.704 | -0.581** | -0.900, -0.246 |
| Nest location | Belowground | 0.183 | -0.523, 1.053 | 0.328 | -0.114, 0.727 |
|  | Aboveground | 0.331 | -0.342, 0.922 | 0.240 | -0.256, 0.566 |
|  | Kleptoparasitic | -0.949** | -1.207, -0.730 | -0.866*** | -1.020, -0.733 |
| Wing length | Continuous (mm) | -0.714** | -1.294, -0.225 | -1.087*** | -1.546, -0.604 |
| Head width | Continuous (mm) | 0.092 | -0.522, 0.612 | 0.181* | 0.092, 0.581 |
| Eye length | Continuous (mm) | 0.977** | 0.558, 1.278 | 1.138*** | 0.955, 1.319 |
| Thorax hair length | Continuous (mm) | -0.713* | -1.045, -0.228 | -0.559*** | -0.749, -0.472 |
| Corbicula setae length | Continuous (mm) | 0.877** | 0.456, 1.264 | 0.529*** | 0.334, 0.803 |
| Corbicula length | Continuous (mm) | -0.301 | -0.527, 0.245 | -0.583** | -0.915, -0.130 |

Taxonomic and phylogenetic nestedness were higher than null model expectations in urban and agricultural landscapes, suggesting high levels of species loss among communities. Functional nestedness (β_sne_) also trended higher than null model expectations, but this was only significant in urban landscapes.

Several traits related to dispersal, nesting, and resource capture contributed to patterns of observed functional beta-diversity (Table 2B). Within urban and agricultural landscapes, bumble bee species were smaller in size, had shorter wing length, were less hairy, but had larger eyes and longer setae on the corbicula than null model expectations based on the regional species pool. Moreover, kleptoparasitic species were less common than null model expectations, whereas the occurrence of species that nest above- or belowground was as expected based on the null model. Several additional traits showed patterns of filtering in urban landscapes only: bumble bee species had shorter corbicula and wider heads, and species with short tongues were more common, while species with long tongues were less common. Although smaller bumble bee species were more common in urban landscapes, these species had a greater range in the body size of the worker caste.

### Anthropogenic species pools to local greenspaces

Based on the urban and agricultural species pools, patterns of observed beta-diversity among local greenspaces were more nuanced. In urban and agricultural landscapes, total taxonomic beta-diversity was as expected based on the null model, while observed total phylogenetic beta-diversity was higher than null model expectations. In contrast, functional beta-diversity showed a different response based on the anthropogenic species pools (Fig. 3; Table 3A). Bumble bee communities sampled in urban and agricultural landscapes contained species that were more phylogenetically distinct than expected based on the landscape-level anthropogenic species pool comparisons, and these patterns were driven by both high turnover and nestedness.

**Figure 3.**
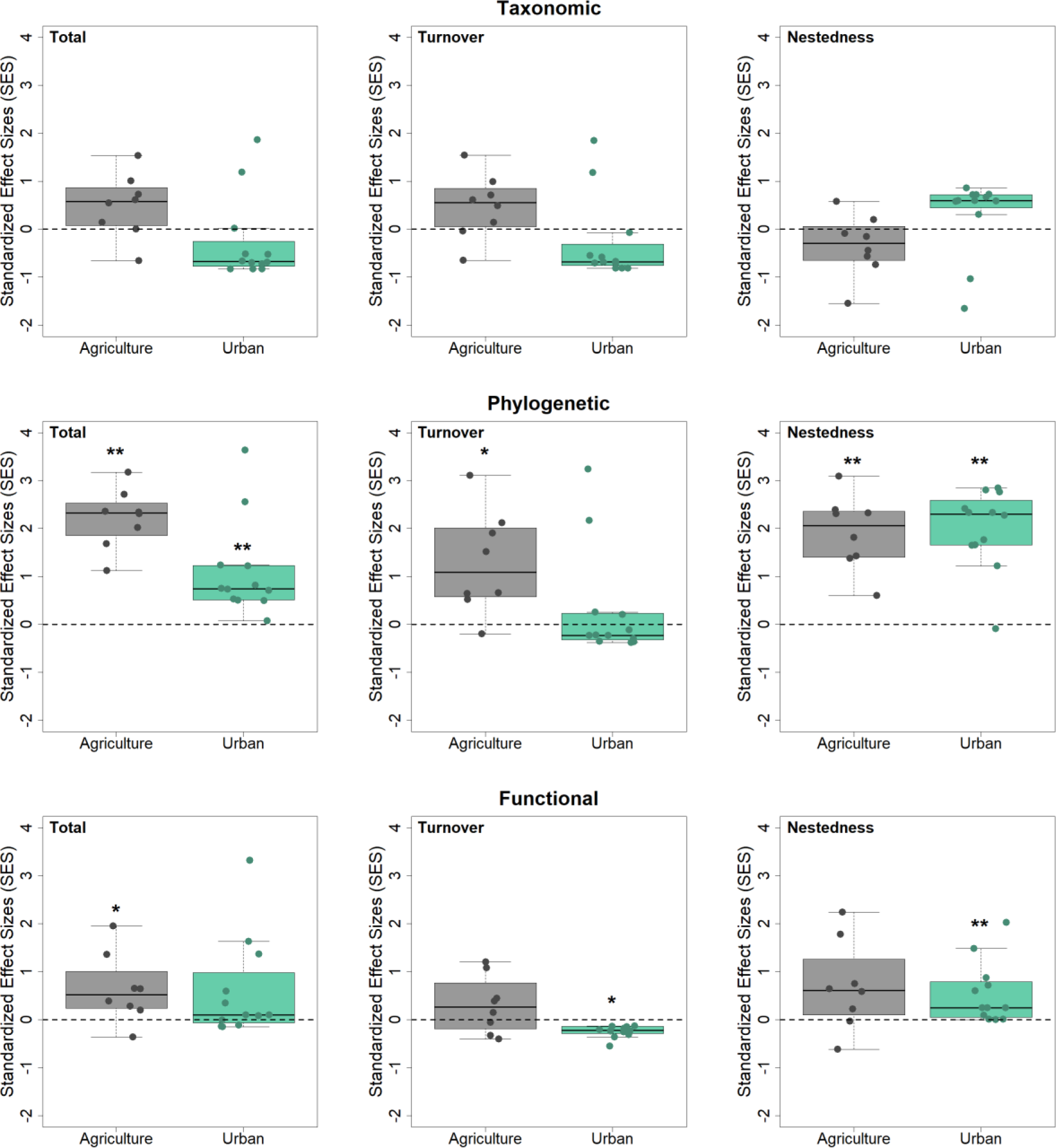
Standardized effect sizes (SES) of observed bumble bee taxonomic (top), phylogenetic (middle), and functional (bottom) beta-diversity in greenspaces embedded in agricultural (left) and urban (right) landscapes surrounding Madison, Wisconsin, USA. Observed values were compared to null model expectations based on the anthropogenic (i.e., urban or agricultural) pool of nine species (Fig. 1B, Local- level Filter). Positive SES values indicate observed beta-diversity metrics were higher than null model expectations, while negative SES values indicate observed beta-diversity metrics were lower than null model expectations. Pairwise Sorensen dissimilarity (β_sor_) matrices were calculated for taxonomic, phylogenetic, and functional beta-diversity. Total beta-diversity for each was decomposed into a turnover component (β_sim_; reflects replacement) and a nestedness component (β_sne_; reflects loss or gain) (i.e., β_sor_ = β_sim_ + β_sne_). Wilcoxon rank sum tests were used to assess whether SES values differed from null model expectations. * P<0.05, ** P<0.01, ***P<0.001, otherwise results were not significant at P>0.05.

**Figure 4.**
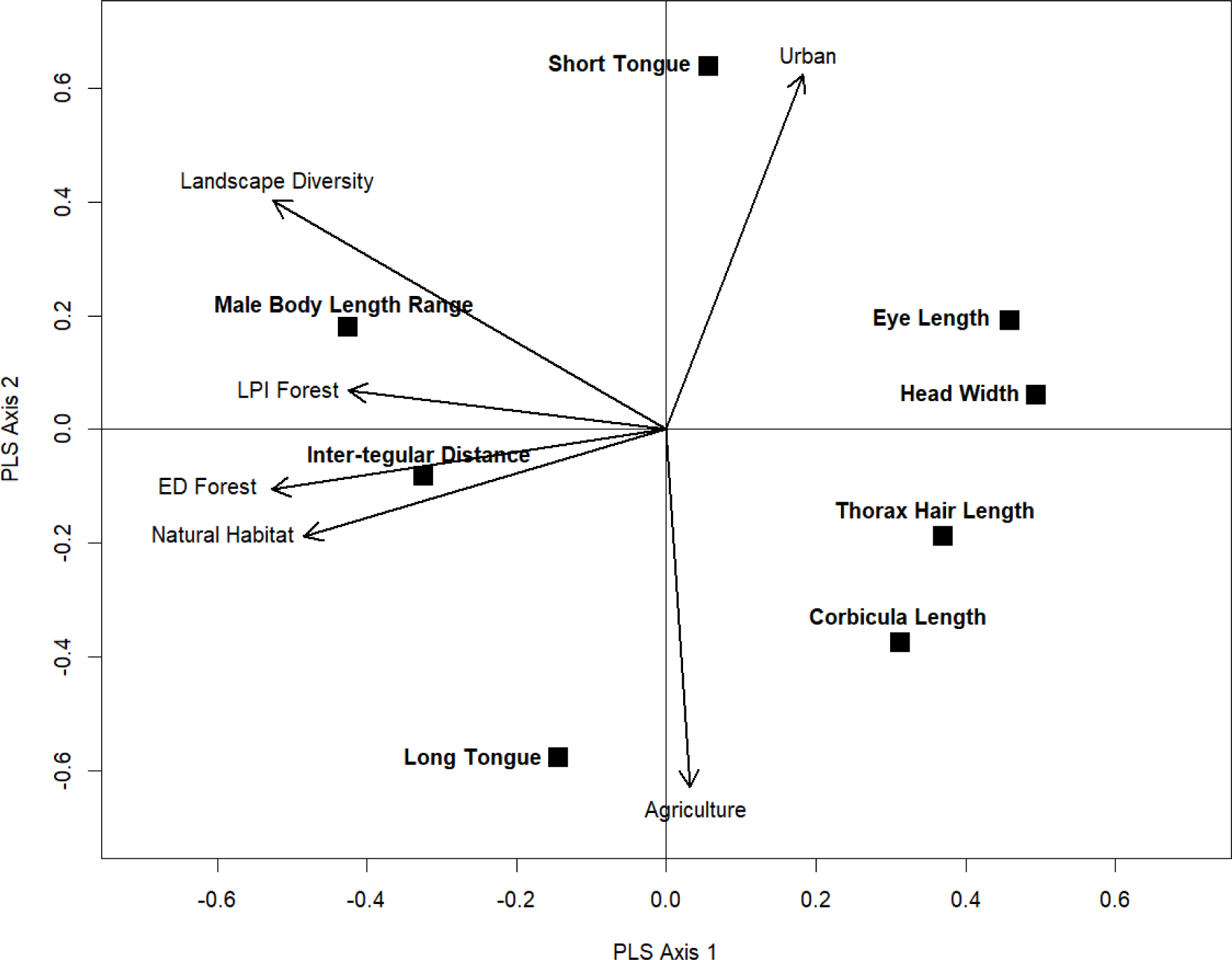
Reduced partial least squares canonical analysis (PLSCA) plot for bumble bee community- weighted means (CWM) for each trait and landscape variables measured at 1.5 km. Total variance explained by axes 1 and 2 in the reduced model for CWMs and landscape variables was 89.7% and 78.6%, respectively. The strength and direction of relationships in PLSCA are determined by relative distance, with closer variables being positively correlated to one another. Bumble bee trait variables are black squares. Landscape variables are indicated by arrows. Configuration and connectivity metrics for Forests were largest patch index (LPI Forest), edge density (ED Forest), and Euclidean nearest neighbor distance (ENN Forest).

**Table 3.** Standardized effect sizes (SES) of observed bumble bee taxonomic, phylogenetic, and functional beta-diversity as well as functional alpha-diversity and community-weighted means for each trait in greenspaces embedded in urban and agricultural landscapes surrounding Madison, Wisconsin, USA. Observed values are compared to null model expectations based on the anthropogenic (i.e., urban or agricultural) pool of nine species (Fig. 1B, Local-level Filter). Positive SES values indicate observed metrics were higher than average null model expectations, while negative SES values indicate observed metrics were lower than average null model expectations. Wilcoxon rank sum tests were used to assess whether SES values differed from null model expectations. * P<0.05, ** P<0.01, ***P<0.001, otherwise results were not significant at P>0.05.

| Metric | Category/Index | Agriculture |  | Urban |  |
| --- | --- | --- | --- | --- | --- |
|  |  | SES<br>(pseudo)median | 95% CI | SES<br>(pseudo)median | 95% CI |
| A) |  |  |  |  |  |
| Taxonomic beta-diversity | Total | 0.522 | -0.058, 1.070 | -0.607 | -0.762, 0.515 |
|  | Turnover | 0.479 | -0.083, 1.077 | -0.628 | -0.756, 0.515 |
|  | Nestedness | -0.312 | -0.998, 0.212 | 0.588 | -0.366, 0.713 |
| Phylogenetic beta-diversity | Total | **2.253\*\*** | 1.680, 2.755 | **0.860\*\*** | 0.523, 1.887 |
|  | Turnover | **1.277\*** | 0.231, 2.308 | -0.067 | -0.296, 1.209 |
|  | Nestedness | **1.878\*\*** | 1.205, 2.696 | **2.049\*\*** | 1.435, 2.558 |
| Functional beta-diversity | Total | **0.519\*** | 0.015, 1.302 | 0.315 | -0.023, 1.499 |
|  | Turnover | 0.315 | -0.231, 0.822 | **-0.220\*** | -0.314, -0.133 |
|  | Nestedness | 0.624 | -0.033, 1.493 | **0.429\*\*** | 0.125, 1.017 |
| B) |  |  |  |  |  |
| Body length | Inter-tegular distance | -0.210 | -1.199, 0.611 | **-0.574\*\*** | -0.942, -0.233 |
| Body length variance | Queen | -0.310 | -0.988, 0.401 | -0.360 | -0.900, 0.104 |
|  | Male | -0.081 | -0.945, 0.965 | -0.239 | -0.388, 0.088 |
|  | Worker | 0.272 | -0.810, 1.078 | **0.503\*** | 0.110, 1.046 |
| Tongue length | Short | -0.505 | -1.455, 0.478 | 0.189 | -0.078, 0.510 |
|  | Medium | **1.079\*\*** | 0.681, 1.465 | **1.046\*\*\*** | 0.791, 1.319 |
|  | Long | -0.350 | -1.263, 0.475 | **-1.148\*\*\*** | -1.463, -0.781 |
| Nest location | Belowground | -0.257 | -1.00, 0.863 | -0.113 | -0.570, 0.348 |
|  | Aboveground | 0.257 | -0.863, 1.008 | 0.113 | -0.348, 0.570 |
| Wing length | Continuous (mm) | -0.528 | -1.287, 0.110 | **-0.929\*\*** | -1.449, -0.329 |
| Head width | Continuous (mm) | 0.043 | -0.814, 0.643 | **0.181\*** | 0.072, 0.724 |
| Eye length | Continuous (mm) | **0.901\*** | 0.106, 1.248 | **1.109\*\*\*** | 0.797, 1.348 |
| Thorax hair length | Continuous (mm) | 0.184 | -0.316, 0.733 | **0.235\*** | 0.016, 0.355 |
| Corbicula setae length | Continuous (mm) | 0.238 | -0.529, 1.052 | -0.355 | -0.726, 0.170 |
| Corbicula length | Continuous (mm) | -0.436 | -0.903, 0.203 | **-0.665\*\*** | -0.960, -0.167 |

Functional beta-diversity showed a different response to the anthropogenic species pools. Total functional beta-diversity was higher than null model expectations in agricultural landscapes, suggesting the traits of bumble bee communities at the local level were more dissimilar than expected based on the traits of the anthropogenic species pool (Fig. 3; Table 3A). In urban landscapes, the turnover in traits among local greenspaces was lower than expected, while the loss of traits among communities was higher than expected based on the null model.

Several traits related to dispersal and resource capture contributed to these patterns of observed functional beta-diversity among local greenspaces in urban and agricultural landscapes (Table 3B).

Within urban landscapes, bumble bee species were smaller in size and had shorter wing length and corbiculae, but had wider heads, larger eyes, and were more hairy than null model expectations based on the urban species pool. Moreover, species with medium tongue length were more common, while species with long tongue length were less common than expected in urban landscapes. Although smaller bumble bee species were more common in urban landscapes, these species had a greater range in the body size of the worker caste. In agricultural landscapes, only two traits differed from null model expectations based on the agricultural species pool. Bumble bee species with medium tongue length and larger eyes were more common than expected in agricultural landscapes.

Bumble bee traits observed in greenspaces based on the local communities were strongly associated with natural habitat and landscape diversity (Fig. 3; Table S2). Landscapes with higher patch diversity, more natural habitat, and larger patches of contiguous forest were positively associated with larger-bodied bumble bee species and species that had a greater range in the body size of the male caste. Landscapes with these same characteristics were negatively associated with bumble bee species that had wider heads, larger eyes, longer corbicula, and longer hair on the thorax. Bumble bee species with short tongues were positively associated with percentage urban, while long tongue species were positively associated with percentage agriculture. Long tongue species also were positively associated with percentage natural habitat and forest edge habitat (ED).

## DISCUSSION

This study investigated whether anthropogenic landscapes dominated by urban or agricultural land cover resulted in biotic homogenization of bumble bee communities in greenspaces along an urban-agricultural gradient measured across three different dimensions of diversity. Our results revealed that only nine of the 13 species from the regional pool were collected in greenspaces along the urban-agricultural gradient, but all nine of these species were found in both urban and agricultural landscapes, indicating identical anthropogenic species pools. However, we found evidence of taxonomic, phylogenetic, and functional homogenization among bumble bee communities relative to the regional species pool (Fig. 1A), regardless of the dominant land cover type (i.e., agricultural or urban). When we assessed filtering from the anthropogenic species pools to local greenspaces (Fig. 1B), we found nuanced differences among land cover types, wherein agricultural landscapes supported more diverse bumble bee communities than expected while urban landscapes continued to show signals of homogenization. Lastly, greenspaces surrounded by more natural habitat supported larger-bodied bumble bee species.

### Assembly of bumble bee communities across spatial scales

Analyses of beta-diversity from the regional species pool to the anthropogenic species pools revealed taxonomic, phylogenetic, and functional homogenization of bumble bees. At the regional scale, anthropogenic landscapes dominated by urban and agricultural land cover acted as strong environmental filters that broadly selected for a subset of functionally similar and phylogenetically related species, which increased the similarity (i.e., reduced beta-diversity) of bumble bee communities among greenspaces. The identical urban and agricultural pools of nine bumble bee species suggests human-dominated landscapes may favor a subset of species regardless of the dominant land cover type and associated resources, shifting communities such that they become dominated by generalist, widespread species. Of the nine species in the anthropogenic species pools, three phylogenetically clustered species (*B. impatiens*, *B. vagans*, and *B. bimaculatus*; Cameron et al. 2007) occurred frequently among greenspaces in both landscapes, contributing strongly to observed patterns of diversity. Consistent with our findings, strong environmental filters at regional scales have been found to dominate processes of insect community assembly over local effects in human-dominated ecosystems (Gámez-Virués et al. 2015, Sol et al. 2017, Fournier et al. 2020, Perry et al. 2020). Although previous research has identified urban and agricultural landscapes individually as environmental filters that lead to biotic homogenization of other taxonomic groups (Ekroos et al. 2010, Karp et al. 2012, Gámez-Virués et al. 2015, Sol et al. 2017), this study revealed that patterns of filtering for bumble bees were largely consistent among these land cover types at large spatial scales.

Compared to the regional scale, patterns of bumble bee community assembly from the urban and agricultural species pools to local greenspaces indicated a reduction in strength of filtering processes as well as nuanced differences among land cover types. Although both anthropogenic pools contained the same bumble bee species, greenspaces in agricultural landscapes supported more functionally diverse communities than expected, while urban landscapes continued to show signals of biotic homogenization. These opposing patterns suggest that variation among greenspaces in agricultural landscapes resulted in higher beta-diversity due to greater species turnover among sites. Agricultural and urban landscapes differ based on a variety of environmental conditions (e.g., amount of impervious and built surfaces, amount of natural habitat, temperature regimes), and these context dependencies can influence the quantity and quality of floral and nesting resources available for bumble bees. Although we did not assess local variables (i.e., floral abundance and richness) in this study, a by-species occupancy analysis by Nunes et al. (X) found that local availability of floral resources was more important than landscape features for limiting the occupancy of bumble bee species in greenspaces across the same urban-agricultural gradient investigated in this study. The finding of higher beta-diversity in bumble bees among agricultural sites, but not across urban greenspaces, could be due to differences in floral resources or other habitat characteristics that are inherently different among agricultural and urban areas.

### Functional diversity of bumble bees in anthropogenic landscapes

The ecological and life history traits of bumble bees influenced how anthropogenic landscapes selectively favored species from the larger regional pool. Traits related to dispersal, nesting habitat, thermoregulatory capabilities, and resource capture emerged as important in determining which species were more common in anthropogenic landscapes. Observed bumble bee communities were characterized by species that were smaller, had shorter wings, larger eyes, reduced hairiness of the thorax, and longer setae on the corbicula (pollen-carrying hind legs). Shorter hair length in anthropogenic environments would be expected if these areas have higher temperatures (e.g., urban heat islands), since longer hair is hypothesized to contribute to thermal insulation (Peters et al. 2016), and hairiness of bees has been documented to decrease with decreasing elevation (i.e., increasing temperatures) (Osorio-Canadas et al. 2022). Although the relationship between pollen load capacity and setae length on the corbicula warrants further study, the length of these setae was related to thorax width in *B. terrestris* (Goulson et al. 2002) and hypothesized to allow smaller workers to carry greater pollen loads relative to their mass (Peat et al. 2005b). This trait may be particularly beneficial in landscapes where floral resource availability is temporally inconsistent and/or spatially patchy such as urban and agricultural ecosystems (Hemberger et al. 2023). While our results are consistent with others that have shown pollinator communities in urban ecosystems select for smaller species (Banaszak-Cibicka and Żmihorski 2012, Eggenberger et al. 2019), other studies found the opposite pattern with larger species more likely to be found in urban areas (Merckx et al. 2018, Fournier et al. 2020, Theodorou et al. 2021).

It is notable that four bumble bee species from the regional pool (*B. citrinus*, *B. pensylvanicus*, *B. perplexus*, and *B. ternarius*) were not observed during the study. These four species had traits that were less common among bumble bee communities than expected by chance, such as having large body size and being kleptoparasites. Kleptoparasitic species are considered indicator taxa for the health and stability of bee communities because they require the presence of their hosts to survive and therefore, are indicative of a higher trophic level within the community (Sheffield et al. 2013).

Several traits were identified as being more consistently found in urban landscapes relative to null model expectations, suggesting urbanization may represent a stronger environmental filter than agriculture for bumble bees. In urban landscapes, bumble bee species with wider heads, shorter corbicula, and short tongues were more common, while long tongue species were less common. These traits are related to resource capture and foraging preference. Tongue length reflects foraging preferences and determines accessibility of nectar based on corolla depth (Goulson et al. 2005). Compared to specialized long tongue species, bumble bees with short tongues can exploit a broad range of floral resources (Goulson et al. 2005, Huang et al. 2015), which may be advantageous in environments with more homogenous floral resources. Corbicula length can influence bumble bee foraging range and duration (Eggenberger et al. 2019), with smaller corbicula resulting in a lower pollen load capacity. Moreover, species that had higher variability in the body size of the worker caste were more common in urban landscapes. Adult body size is related to the amount of food received during larval development (Sutcliffe and Plowright 1988), allocation and provision strategies within the colony (Schmid-Hempel and Schmid-Hempel 1998, Cueva del Castillo et al. 2015), and nesting ecology (Roulston and Cane 2000). Increased variability in body size among foragers within a colony may increase access to a greater diversity of floral resources within the urban environment as individuals vary in their foraging distance and efficiency (Spaethe and Weidenmüller 2002, Goulson et al. 2002, Peat et al. 2005a, Greenleaf et al. 2007) and flower handing time (Peat et al. 2005b). This colony level phenotypic plasticity may be a successful strategy in urban areas where floral resources may be diverse but their availability and spatial distribution within the environment is patchy (Eggenberger et al. 2019).

### Importance of natural habitat in anthropogenic landscapes

Several bumble bee traits were associated with the amount and connectivity of natural habitat and landscape diversity. Within this region, natural habitat was largely dominated by forest and wetland ecosystems. Although all greenspaces sampled in this study were dominated (>50%) by either urban or agricultural land cover, greenspaces varied (range <1% to 47.5%) in the amount of natural forest and wetland habitat that was present within the surrounding 1.5 km landscape. Greenspaces surrounded by more diverse land cover types and more natural habitat, including larger patches of contiguous forest and more forest edge habitat, supported on average larger bumble bee species, species with greater variability in the body length of the male caste, and species with long tongues. Body size has been linked to dispersal capacity in bees where larger species are capable of foraging at greater distances than smaller species (Greenleaf et al. 2007). Therefore, larger bees may be better able to access fragmented patches of forest in anthropogenic landscapes. Recent studies have highlighted the value of forests as complimentary habitat for bumble bee biology and population persistence. Forests contain important spring ephemeral forage plants for bumble bees, and this resource pulse overlaps with queen activity and colony establishment (Mola et al. 2021b). Forests also provide nesting habitat for bumble bee colonies during the growing season and overwintering habitat for queens (Mola et al. 2021a). Our findings suggest that the amount and composition of natural habitat surrounding greenspaces selects for bumble bee species that have a different set of traits, even if these natural habitat types are not the dominant land cover. The amount of natural habitat surrounding homogenous or resource poor environments such as crop monocultures or urban areas can modify bee communities by supporting higher turnover and overall diversity of species and traits in the landscape. High species and trait diversity could ultimately benefit pollination services, with benefits being complimentary to local resource availability and site management (Kremen et al. 2007, Kennedy et al. 2013).

## Conclusions

Community assembly is a multi-scale process determined by a series of hierarchical filters that cumulatively influence which species from the regional species pool can colonize and establish to form local communities (HilleRisLambers et al. 2012, Aronson et al. 2016, Perry et al. 2020). To support pollinator conservation in human-dominated landscapes, there is a need to understand the drivers of community assembly at large spatial scales. We documented evidence of taxonomic, phylogenetic, and functional homogenization among bumble bee communities at the regional scale, and these patterns were similar among urban and agricultural landscapes, indicating that landscape level filters selected for a subset of functionally similar and phylogenetically related species that formed anthropogenic species pools. Although urban and agricultural species pools contained the same bumble bee species, nuanced patterns of assembly were observed among greenspaces at local scales, as among site variation in abiotic and/or biotic filters acted within these landscape types to influence the occurrence of bumble bee species.

Biotic homogenization (i.e., low turnover and high nestedness) as was observed in this study of bumble bee communities may impact the ecosystem services and function provided by this important group of pollinators at regional scales. For example, the relative importance and strength of species turnover for pollination services was found to increase with increasing spatial scales such that commercial crop fields distributed across a 3,700 km^2^ landscape required nearly 80 bee species to achieve a 75% threshold level of pollination (Winfree et al. 2018). Although this study focused on only one group of insect pollinators, the importance of bumble bee contributions to pollination services has been documented in natural, agricultural, and urban settings (Memmott et al. 2004, Goulson et al. 2008, Matteson and Langellotto 2009). Likewise, the parallel loss of bumble bee diversity across urban and agricultural landscapes may have compounded effects on pollination services, as well as the capacity for these communities to respond to future environmental change. Our findings support a landscape-level approach to biodiversity conservation that promotes diversifying landscapes to support pollinator populations and ongoing conservation initiatives. To achieve this management strategy, interdisciplinary collaboration and cooperation among multiple partners is required to implement conservation measures and succeed at creating multi-functional landscapes.

## ACKNOWLEDGEMENTS

This research was supported by a USDA National Institute of Food and Agriculture grant (2019-67013- 29298). We thank Jade Kochanski, Murilo Alvez-Zacareli, and Sammy Herdman for assistance with field sampling, and the landowners and land managers who provided access to private lands for sampling. We thank Heather Hines for sharing the bumble bee phylogeny that was integral to this research.

