## Appendix S1 for "Homogenization of taxonomic, phylogenetic, and functional characteristics of bumble bee communities at regional scales in anthropogenic landscapes"

**Figure S1.** Pearson correlation analyses were used to assess the relationships among bumble bee traits. Traits that were highly correlated were removed from further analyses. The traits shown below are: Body length range for queen (qbl_var), male (mbl_var), and worker (wbl_var) castes; tongue length (tl); nest location (nestl); inter-tegular distance (it); wing marginal cell length (rwingmc); wing width (rwingw); wing length (rwingl); head width (rheadw); eye width (reyew); eye length (reyel); scape length (rscapel); length of hair on the thorax (thairl); length of setae on the posterior edge of the corbicula (bsetael); corbicula length (rbtl); and corbicula width (rbtw). Wing marginal cell length, wing width, eye width, corbicula width, and scape length had correlation coefficients of 0.60 or higher with other traits and were removed.

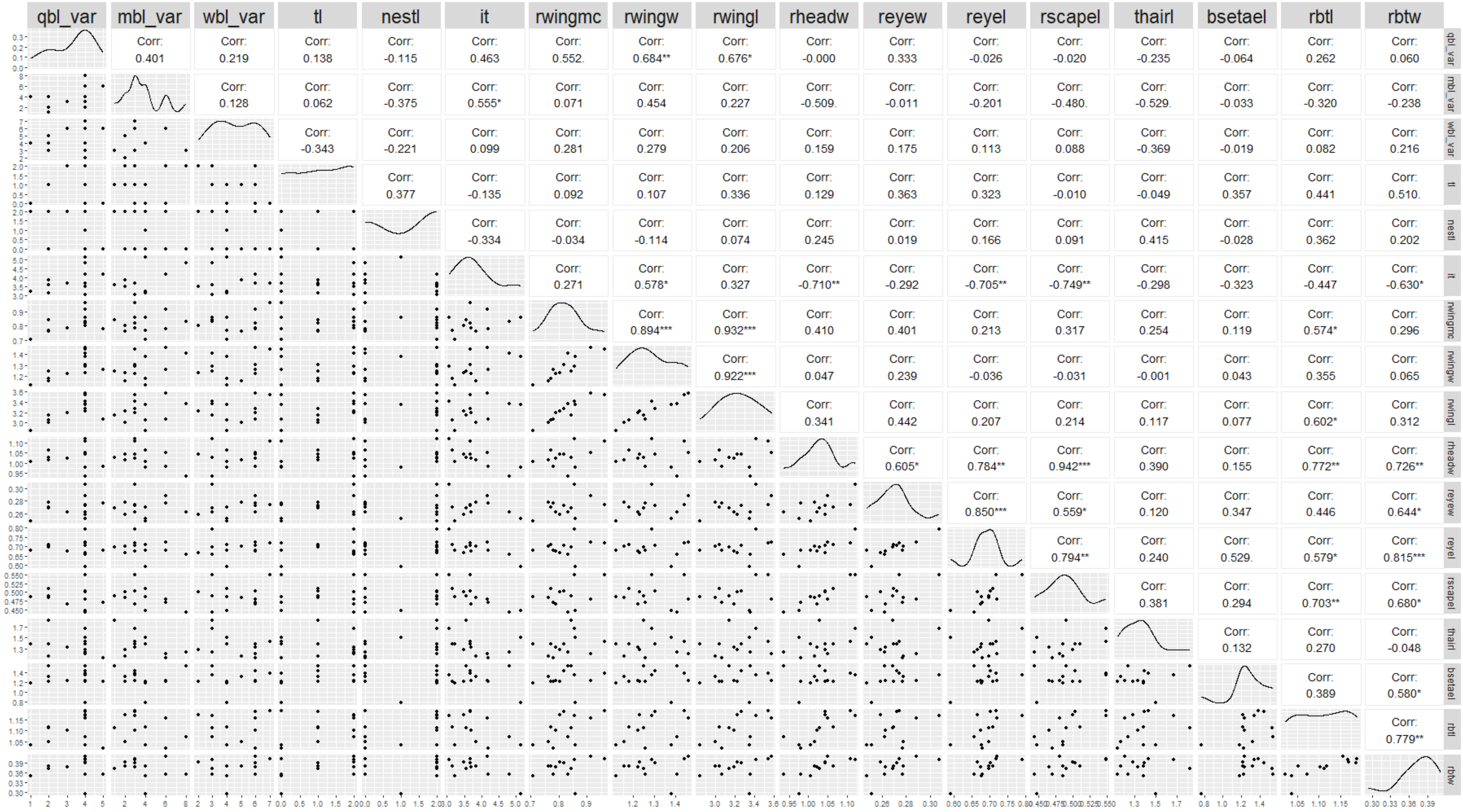
qbl_var
mbl_var
wbl_var
tl
nestl
it
rwingmc
rwingw
rwingl
rheadw
reyew
reyel
rscapel
thairl
bsetael
rbtl
rbtw
Corr: 0.401
Corr: 0.219
Corr: 0.138
Corr: -0.115
Corr: 0.463
Corr: 0.552.
Corr: 0.684**
Corr: 0.676*
Corr: -0.000
Corr: 0.333
Corr: -0.026
Corr: -0.020
Corr: -0.235
Corr: -0.064
Corr: 0.262
Corr: 0.060
Corr: 0.128
Corr: 0.062
Corr: -0.375
Corr: 0.555*
Corr: 0.071
Corr: 0.454
Corr: 0.227
Corr: -0.509.
Corr: -0.011
Corr: -0.201
Corr: -0.480.
Corr: -0.529.
Corr: -0.033
Corr: -0.320
Corr: -0.238
Corr: -0.343
Corr: -0.221
Corr: 0.099
Corr: 0.281
Corr: 0.279
Corr: 0.206
Corr: 0.159
Corr: 0.175
Corr: 0.113
Corr: 0.088
Corr: -0.369
Corr: -0.019
Corr: 0.082
Corr: 0.216
Corr: 0.377
Corr: -0.135
Corr: 0.092
Corr: 0.107
Corr: 0.336
Corr: 0.129
Corr: 0.363
Corr: 0.323
Corr: -0.010
Corr: -0.049
Corr: 0.357
Corr: 0.441
Corr: 0.510.
Corr: -0.334
Corr: -0.034
Corr: -0.114
Corr: 0.074
Corr: 0.245
Corr: 0.019
Corr: 0.166
Corr: 0.091
Corr: 0.415
Corr: -0.028
Corr: 0.362
Corr: 0.202
Corr: 0.271
Corr: 0.578*
Corr: 0.327
Corr: -0.710**
Corr: -0.292
Corr: -0.705**
Corr: -0.749**
Corr: -0.298
Corr: -0.323
Corr: -0.447
Corr: -0.630*
Corr: 0.894***
Corr: 0.932***
Corr: 0.410
Corr: 0.401
Corr: 0.213
Corr: 0.317
Corr: 0.254
Corr: 0.119
Corr: 0.574*
Corr: 0.296
Corr: 0.922***
Corr: 0.047
Corr: 0.239
Corr: -0.036
Corr: -0.031
Corr: -0.001
Corr: 0.043
Corr: 0.355
Corr: 0.065
Corr: 0.341
Corr: 0.442
Corr: 0.207
Corr: 0.214
Corr: 0.117
Corr: 0.077
Corr: 0.602*
Corr: 0.312
Corr: 0.605*
Corr: 0.784**
Corr: 0.942***
Corr: 0.390
Corr: 0.155
Corr: 0.772**
Corr: 0.726**
Corr: 0.850***
Corr: 0.559*
Corr: 0.120
Corr: 0.347
Corr: 0.446
Corr: 0.644*
Corr: 0.794**
Corr: 0.240
Corr: 0.529.
Corr: 0.579*
Corr: 0.815***
Corr: 0.381
Corr: 0.294
Corr: 0.703**
Corr: 0.680*
Corr: 0.132
Corr: 0.270
Corr: -0.048
Corr: 0.389
Corr: 0.580*
Corr: 0.779**

**Table S1**. Partial least squares canonical analysis (PLSCA) correlation coefficients for bumble bee community-weighted means (CWM) for each trait and landscape variables at 1.5 km. Variables with a correlation coefficient higher than 0.30 or lower than -0.30 were retained for a reduced PLSCA model and relevance network analysis. In the full model, total variance explained by axes 1 and 2 was 81.9% and 51.7%, respectively. In the reduced model, total variance explained was 89.7% by axis 1 and 78.6% by axis 2.

| **Metric** | **Category/Index** | **Full Model** | | **Reduced Model** | |
|---|---|---|---|---|---|
| | | **PLS Axis 1** | **PLS Axis 2** | **PLS Axis 1** | **PLS Axis 2** |
| *Bumble bee CWMs* | | | | | |
| Body length | Inter-tegular distance | -0.30 | -0.05 | -0.32 | -0.08 |
| Body length variance | Queen | 0.03 | -0.04 | | |
| | Male | -0.40 | 0.17 | -0.42 | 0.18 |
| | Worker | 0.27 | 0.17 | | |
| Tongue length | Short | 0.03 | 0.55 | 0.05 | 0.64 |
| | Medium | 0.20 | -0.14 | | |
| | Long | -0.11 | -0.49 | -0.14 | -0.57 |
| Nest location | Belowground | -0.06 | 0.25 | | |
| | Aboveground | 0.06 | -0.25 | | |
| Wing length | Continuous (mm) | 0.11 | -0.20 | | |
| Head width | Continuous (mm) | 0.45 | 0.45 | 0.49 | 0.06 |
| Eye length | Continuous (mm) | 0.42 | 0.15 | 0.45 | 0.19 |
| Thorax hair length | Continuous (mm) | 0.32 | -0.15 | 0.36 | -0.18 |
| Corbicula setae length | Continuous (mm) | -0.16 | -0.20 | | |
| Corbicula length | Continuous (mm) | 0.27 | -0.31 | 0.31 | -0.37 |
| *Landscape variables* | | | | | |
| Landscape diversity | Simpson diversity index | -0.50 | 0.41 | -0.52 | 0.40 |
| Percentage agriculture | Percentage at 1.5 km | 0.03 | -0.64 | 0.03 | -0.63 |
| Percentage urban | Percentage at 1.5 km | 0.17 | 0.59 | 0.18 | 0.62 |
| Percentage natural habitat | Percentage at 1.5 km | -0.47 | -0.14 | -0.48 | -0.18 |
| LPI forest | Largest patch index | -0.41 | 0.10 | -0.42 | 0.06 |
| ED forest | Mean edge density | -0.51 | -0.08 | -0.52 | -0.10 |
| ENN forest | Euclidean nearest neighbor | 0.23 | -0.15 | | |

**Table S2.** Pairwise similarity matrix calculated from relevance network analyses following reduced partial least squares canonical analyses (PLSCA) for bumble bee community-weighted means (CWM) for each trait and landscape variables at 1.5 km. Similarity values are calculated by summing the correlations between the individual variables and each of the latent components from the PLSCA model. These similarity values approximate a Pearson correlation coefficient. A 0.5 similarity value threshold was used for the relevance network analysis, and variable associations that met this threshold are shown. Landscape configuration and connectivity metrics for class forest were largest patch index (LPI Forest), edge density (ED Forest) and Euclidean nearest neighbor distance (ENN Forest).

| Metric | Category | Landscape Diversity | Percentage Agriculture | Percentage Urban | Percentage Natural Habitat | LPI Forest | ED Forest | ENN Forest |
|---|---|---|---|---|---|---|---|---|
| Body length | Inter-tegular distance | 0.51 | | | 0.67 | 0.65 | 0.68 | |
| Body length variance | Queen | | | | | | | |
| | Male | 0.81 | | | 0.65 | 0.70 | 0.69 | |
| | Worker | | | | | | | |
| Tongue length | Short | | -0.92 | 0.93 | | | | |
| | Medium | | | | | | | |
| | Long | | 0.84 | -0.95 | 0.64 | | 0.58 | |
| Nest location | Belowground | | | | | | | |
| | Aboveground | | | | | | | |
| Wing length | Continuous (mm) | | | | | | | |
| Head width | Continuous (mm) | -0.68 | | | -0.90 | -0.88 | -0.91 | |
| Eye length | Continuous (mm) | -0.53 | | 0.52 | -0.83 | -0.78 | -0.83 | |
| Thorax hair length | Continuous (mm) | -0.59 | | | -0.77 | -0.75 | -0.78 | |
| Corbicula setae length | Continuous (mm) | | | | | | | |
| Corbicula length | Continuous (mm) | -0.74 | | | -0.58 | -0.63 | -0.62 | |
